# Humanized Caffeine-Inducible Systems for Controlling Cellular Functions

**DOI:** 10.1101/2025.06.13.659463

**Authors:** Leo Scheller, Maddalena Elia, Lucia Bonati, Chungwon Kang, Pablo Gainza, Yash Garodia, Pranitha Deshpande, Joseph Schmidt, Tom Enbar, Sandrine Georgeon, Li Tang, Bruno E. Correia

## Abstract

Current cell therapies are limited by the lack of tools for controlling gene expression using humanized systems responsive to non-toxic stimuli. Starting from nanobodies that homodimerize in response to caffeine, we computationally designed inducible heterodimers and humanized the best-performing pairs. We used the resulting caffeine-inducible domains in engineered cytokine receptors for caffeine-inducible STAT3 signaling and in split transcription factors (caff-TFs) containing human-derived zinc-finger proteins. Heterodimerization of split transcription factors drastically enhanced their performance compared to homodimerization. We demonstrate that caff-TFs are compatible with lentiviral and retroviral delivery to Jurkat T-cells and enable inducible expression of therapeutic genes such as chimeric antigen receptors (CARs) in response to caffeine concentrations consistent with normal coffee consumption. By using the common, non-toxic molecule caffeine, and exclusively humanized protein components, these systems promise to be a safer alternative to existing systems and may be used in synthetic biology applications and for safer, more effective cell therapies.

## Introduction

Cell therapies are emerging as a new class of therapeutic approaches for a variety of diseases^1^. However, controlling gene expression and cell signaling in engineered cells remains a critical challenge. For instance, Chimeric Antigen Receptor (CAR)-T cell therapy is highly efficient in treating liquid tumors but requires improved control strategies for successful application to solid tumors^2,3^. Similarly, cell-based expression of cytokines has shown promising results and inducible promoter systems are being investigated to reduce side effects^4–6^.

Numerous synthetic biology tools have been developed based on genetic circuits and receptor systems to regulate signaling and expression levels of effector proteins in response to small molecules^1^. However, the sensor components, such as transcription factors or engineered receptor domains, typically contain proteins of viral, bacterial, plant, or yeast origin. Their potential immunogenicity likely reduces cell persistence and complicates clinical use^7–9^. Additionally, the most common small molecule inducers like tamoxifen or rapamycin, although effective in vitro, have side effect profiles and pharmacokinetic limitations that pose challenges for clinical application^10–13^. Therefore, developing humanized systems for controlling gene expression or receptor activation responsive to non-toxic small molecules could advance cell therapy.

Zinc finger proteins (ZFs) are a major class of human transcription factors and their modular structure can be reassembled to bind new DNA sequences. While plant-based Transcription activator-like effector nucleases (TALENs) and bacterial-based dCas9 have become very popular in synthetic biology due to their high programmability^14,15^, ZFs remain a promising option for gene circuits in cell therapy due to their human origin. Recently, ZFs have been developed that are predicted not to bind to any human genomic region, thus providing safe DNA binding domains for therapeutic applications^16^. However, as the authors point out, for small-molecule controlled transcriptional regulation, their system still is limited by either using plant-based or viral proteins or by the need for small molecules with suboptimal safety and side-effect profiles. Here, we aim to combine the established ZF platform with engineered, humanized domains for transcriptional control by caffeine.

Caffeine is consumed worldwide and has become increasingly established as a regulator for protein interactions in synthetic biology through camelid, caffeine-inducible homodimerising nanobodies^17–21^. Its favorable safety profile and pharmacokinetics make it an attractive candidate for clinical applications^22^. However, using caffeine in cell therapy faces two major challenges: the expected immunogenicity of nonhuman protein domains when expressed by a cell, and the need for heterodimerization in various applications, such as split transcription factors (TFs) or split chimeric antigen receptors (CARs). In both cases, homodimerization would lead to nonfunctional states. We hypothesized that computational design could identify complementary point mutations at the caffeine-bound nanobody interface to favor heterodimerization and disfavor homodimerization.

CAR-T cells with camelid nanobodies (variable heavy domain of heavy chain; VHH) as tumor targeting domains have been observed to be highly effective initially, but persistence in patients is low due to clearance by the immune system^23^. Humanization of proteins in cell therapies, including VHHs, have been shown to enhance cell persistence^24,25^. By now, one CAR-T cell therapy with two humanized VHHs and one humanized VHH-based drug has been approved for clinical use^26,27^, demonstrating that humanized nanobodies are a promising avenue for improving cell therapies.

In this work, we developed humanized caffeine-inducible homodimers and heterodimers to control cell function with caffeine. The caffeine-inducible transcription factors (caff-TFs) and caffeine-inducible cytokine receptors (caff-GEMS) were exclusively based on human-derived or humanized components and were compatible with lentiviral and retroviral delivery to T-cells in single-vector systems. This enabled the inducible expression of transgenes, such as GFP or CARs, in response to caffeine concentrations that are within the range of plasma levels observed after coffee consumption. These systems offer safer alternatives to existing gene regulation systems and hold potential for wide-ranging applications in synthetic biology and cell therapy.

## Results

### Computational Design of Caffeine Induced Heterodimers

We used the structure of an anti-caffeine VHH that homodimerizes in response to caffeine (PDB:6qtl)^21^ as a basis for design. To generate protein modules that heterodimerize, we used positive design to make a protein-protein interface that is compatible between heterodimers and negative design to disfavor homodimerization (**Figure 1a**). These systems could enhance performance of systems where homodimerization leads to nonfunctional protein complexes, hampering system performance. Residues interacting with caffeine were discarded from consideration as they would likely result in a loss of affinity. To search the entire sequence/conformational landscape in a provable way, we developed a faster version of the COMETS algorithm^28^, which we call varCOMETS. The top designs all resulted in charged residues at the interface, with the opposite charges facing each other in the heterodimers and same charges facing each other in the homodimers (**Table S3.**). We chose the best seven heterodimer candidates based on the multistate Rosetta energy to test them experientially.

**Figure 1:**
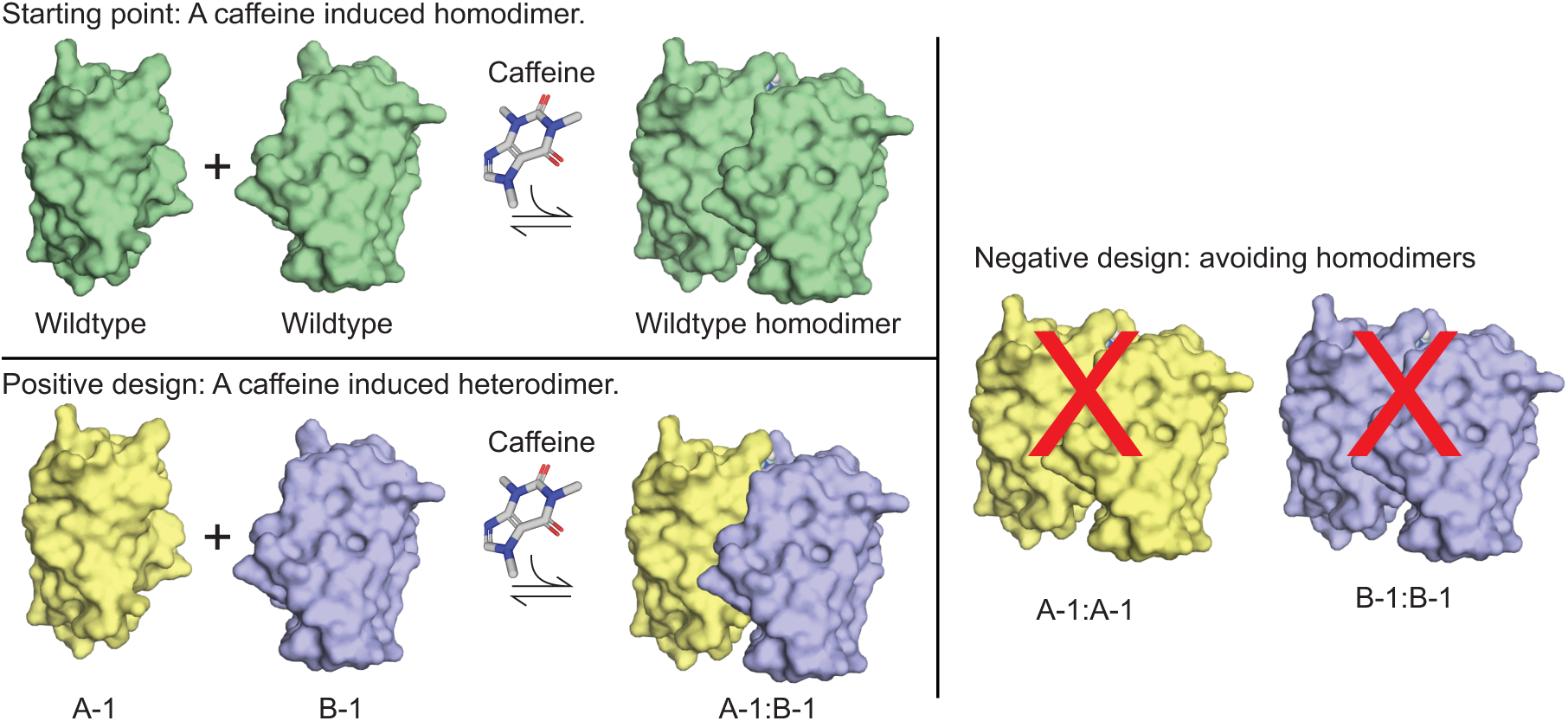
Computational Design Goals. **a,** Schematic of using positive and negative design to create caffeine induced heterodimers. The starting point are camelid nanobodies that homodimerize in response to caffeine. We want to design caffeine induced heterodimerization and avoid homodimerization.

### Heterodimer Selection from Top 7 Designs

To test heterodimerization in response to caffeine, we used the GEMS receptor platform that drives reporter gene expression in response to dimerization (**Figure 1c**). These receptors can function by both homodimerization and heterodimerization and were previously shown to work in combination with caffeine nanobodies^17,29^. We reasoned that such a system enables estimations of fold-differences in caffeine concentrations required for heterodimerization over homodimerization in a cell-based context.

In this system, the VHH is N-terminally fused to a chimeric receptor consisting of an erythropoietin receptor (EpoR) extracellular and transmembrane domain (ECD, TMD) and an interleukin-6 receptor B (IL-6RB) intracellular domain (ICD). Dimerization of the VHHs activates the IL-6RB ICD and leads to STAT3 phosphorylation, which can be quantified with a reporter plasmid consisting of STAT3 binding sites upstream of a minimal promoter, driving expression of the reporter gene secreted nanoluciferase.

We screened for pairs that have a lower EC_50_ in terms of caffeine-induced reporter expression in cells expressing both heterodimer receptor chains compared to the respective single chains only. The difference in EC_50_ between heterodimers and homodimers was greatest for complex 3 with an EC_50_ of 150 nM compared to 2 or 10 µM respectively for the homodimers (**Figure 2, Figure S1, Table S6**). These results indicate that the resulting heterodimeric caffeine nanobodies are suitable for the design of heterodimerization-dependent systems.

**Figure 2:**
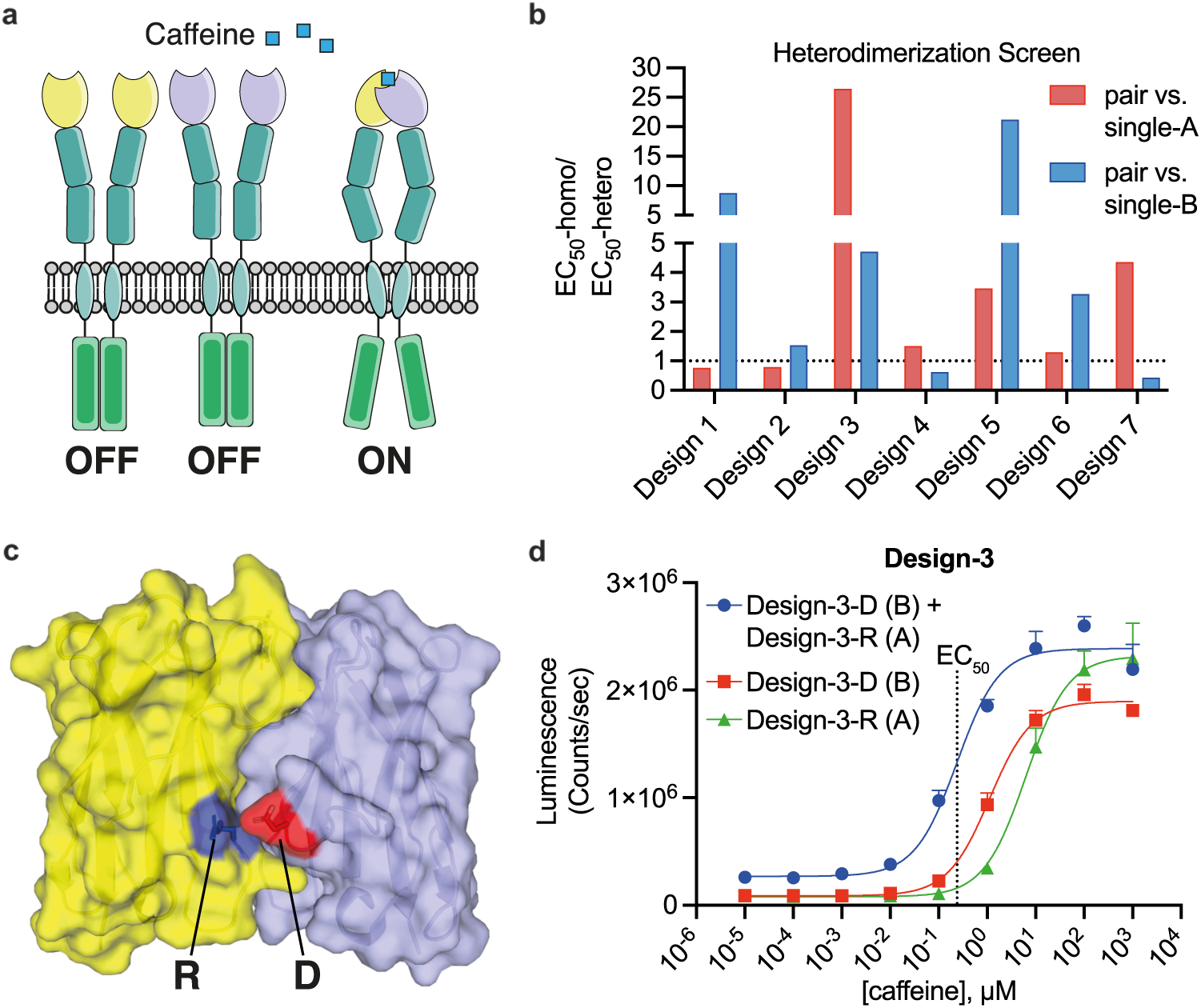
Caffeine Induced Heterodimers. **a,** The GEMS platform was used for assessing designed heterodimer pairs. Heterodimeric receptors were designed to form high sensitivity receptors, whereas homodimers would signal with lower sensitivity in response to caffeine. **b,** Screening results for seven designed heterodimers. The EC50 (the concentration of caffeine required to induce half-maximal reporter expression) for each homodimer is divided by the EC50 of its corresponding heterodimer. Ratios above 1 indicate that the heterodimer is more sensitive to caffeine than the homodimer. Exact values and proteins are listed in table S6 **c,** Structure of the original protein (PDB: 6qtl) with highlighted interacting, charged residues. These residues were inverted in Design 3 (monomer 1: R47D; monomer 2: D64R), leading to unfavorable D:D and R:R homodimers. **d,** dose response curve for Design 3, comparing the caffeine sensitivity of the heterodimer with its corresponding homodimers.

### Homodimer Humanization and Selection

To generate a human-derived GEMS platform, we exchanged the murine EpoR ECD and TMDs with the equivalent human domains. To humanize the VHHs, we introduced all mutations previously described for a maximally humanized VHH^30^ (Humanized-1). Additionally, we introduced the point mutation Y102W, which was reported to increase caffeine sensitivity^18^ (Humanized-2). We next reverted the mutations (G44R, L45R) back to charged residues as in Design-7-R, which may be important for the stability of monomeric heavy chain fragments^31^ (Humanized-3). All resulting receptors still had high basal activity, indicating dimerization in the absence of caffeine (**Figure 3a**).

**Figure 3:**
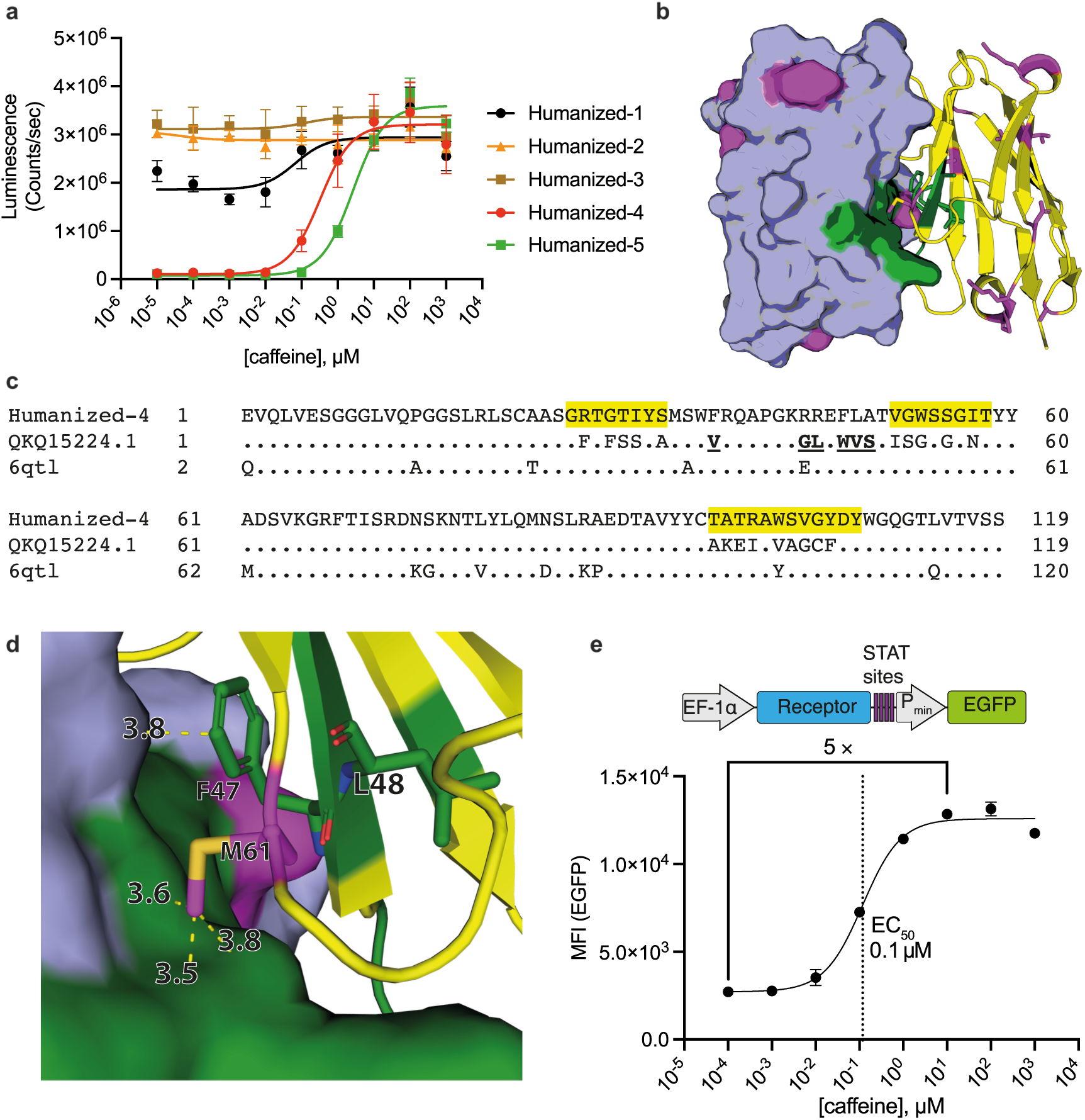
Nanobody Humanization: a, Dose response curves of GEMS containing humanized caffeine nanobodies. **b,** Structure of the original protein (PDB: 6qtl) with residues for humanization shown in magenta and camelid residues shown in green. **c,** Sequence alignment of a the humanized nanobody (Humanized-4) to a top ranked result from a blast search against human proteins (Sbjct; anti-SARS-CoV-2 monoclonal antibody heavy chain variable region, partial [Homo sapiens] Sequence ID: QKQ15224.1) and to the original camelid nanobody (6qtl). The CDRs (IMGT annotation) are highlighted in yellow. Nonhuman residues outside of the CDRs are shown in bold and underlined. **d,** Interface region between the dimerized nanobodies (PDB: 6qtl) with residues F47, L48, M61 and their distances in Å to the other nanobody. Amino acids that were humanized in Humanized-4 are marked in magenta and those that were not humanized in green. **e,** Schematic of an all-in-one lentiviral vector for STAT3 GEMS driving GFP expression and the corresponding dose response in Jurkat cells.

Next, we blast searched the sequence against human proteins and humanized additional residues (Q1E, A35S, M61A; Humanized-4). M61A is a mutation at an interface residue, which we reasoned might reduce background dimerization. By the same reasoning, we additionally tested L48V (Humanized-5) (**Figure 3d**). These final, maximally humanized designs restored very low basal activity with Humanized-4, being the most sensitive with an EC_50_ of 360 nM in the GEMS platform, which is in the same range as the EC_50_ of 210 nM of the wild type camel antibody (**Figure S2**) and well below the typical caffeine plasma concentration of 15 µM after drinking one coffee^22^.

Humanized-4 contains 6 amino acid differences outside of the CDRs (annotated according to the international ImMunoGeneTics; IMGT^32^) to the closest human antibody sequence. By the same metric, the approved humanized Caplacizumab^33^ contains 11 amino acid differences to the closest human antibody sequence (**Figure S3**). All tested mutations are listed in **table S4.** Next, we generated all-in-one lentiviral vectors containing the STAT3 caffeine receptors together with the respective inducible promoter elements for driving EGFP expression. Transduced Jurkat cells showed highly sensitive caffeine-induced EGFP expression (EC_50_ < 1 µM) with a 5-fold increase in reporter expression upon induction compared to the uninduced state. These results demonstrate caffeine-induced cytokine signaling in a human T-cell line by a humanized receptor.

### Heterodimer Humanization, Optimization, and Selection

To generate humanized caffeine induced heterodimers, we introduced the heterodimerization mutations of Design-3 into the Humanized-4 framework. However, in the GEMS, we observed high basal activation in the absence of caffeine. To find the best pair with better signal-to-noise ratio, we compared combinations of nanobodies with mutations at different interface residues (**Figure 4a, table S5**). For some of the most promising constructs, we performed dose response curves and compared the EC_50_s for the heterodimers compared to the respective homodimers and observed greatly improved changes in heterodimer vs homodimer sensitivities (**Figure 4b**). The pairs A3 + B3 had very high sensitivity, but still high background activity. The pair A3 + B6 had a close to 10-fold signal to noise ratio and very high changes in the EC_50_s between homodimers and heterodimers (**Figure 4b,e, table S6**). The heterodimer pairs A2 + B1, A3 + B1 performed with signal to noise ratios of 20 or higher, and the B1 homodimer had an about 7-fold increase in EC_50_ compared to the heterodimers. (**Figure 4 e,f, table S6**). These results show that several heterodimer pairs are promising candidates for extracellular applications. While the homodimerizing variants are sufficient for the GEMS, other split receptor systems or the reconstitution of split proteins rely on heterodimerization and could benefit from these variants.

**Figure 4:**
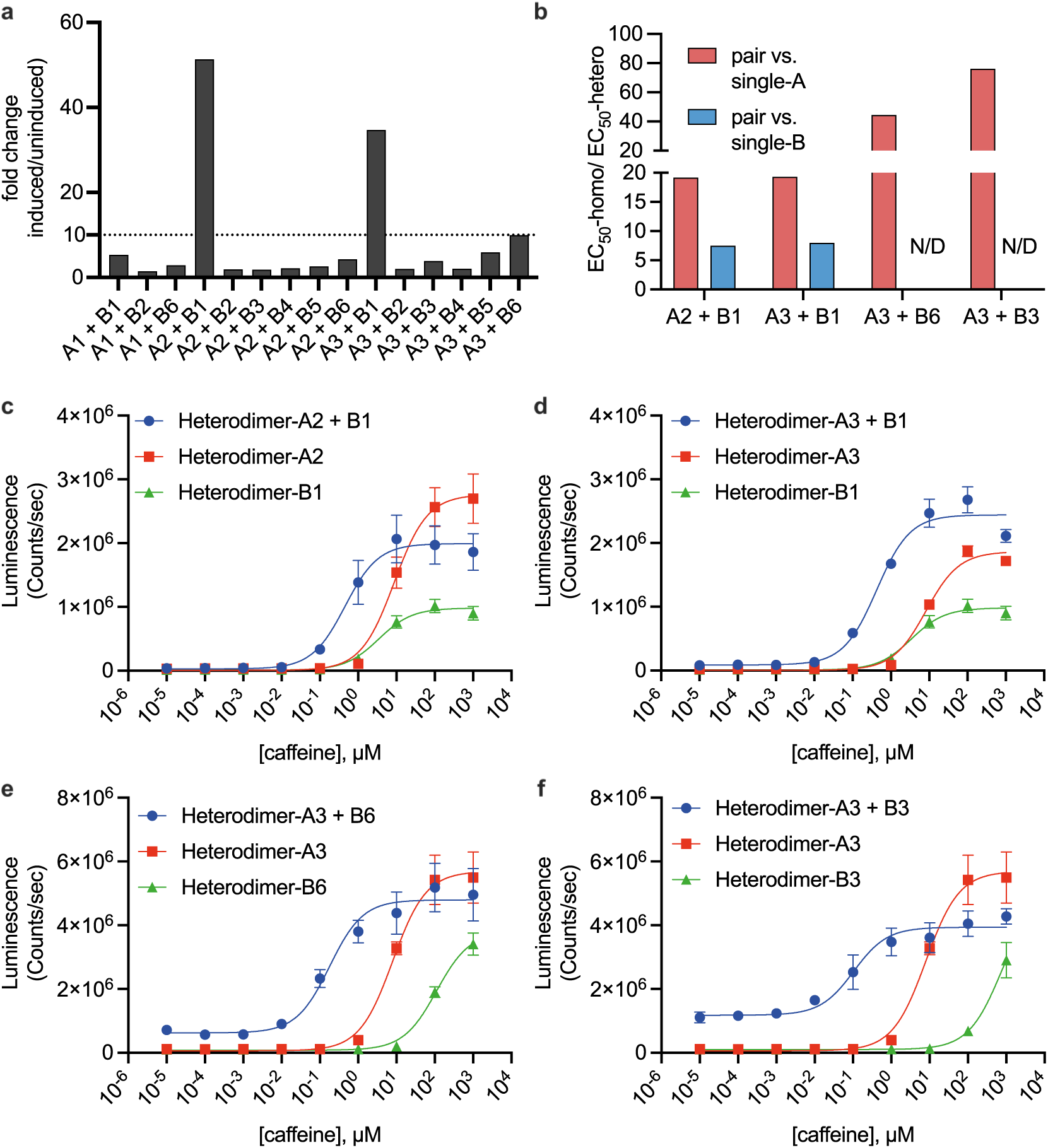
Humanized Heterodimer Selection. **a,** Fold change of reporter activity for induced/uninduced (100 µM caffeine/ no caffeine) of the different heterodimer combinations. **b,** Sensitivity differences of heterodimer vs. individual homodimers by calculating EC50 of each homodimer divided of EC50 by the complex. Smaller EC50 correspond to higher sensitivity, therefore large numbers correspond to bigger differences between heterodimer and homodimer sensitivities. As B3 and B6 do not reach a stable plateau at the highest caffeine concentration, the fold change is not defined, but can be assumed to be the high. **c, d, e, f,** dose response curves for the best performing heterodimers in the GEMS platform. Values in the dose response curves are the mean ± s.d. of n = 3 biological replicates.

### Caffeine-Inducible Control of Zinc Finger Transcription Factors

To generate caffeine inducible transcription factors (caff-TFs), we fused the humanized caffeine nanobody heterodimers to zinc finger domains that were derived from human proteins or to the transactivator domain (TAD) of human RELA/p65. The zinc finger domains were designed to bind to DNA sequences that are not present in the human genome^16^ **(Figure 5a)**. Two of the tested pairs (ZF-B3 together with either A2-TAD or A3-TAD) performed very similarly with EC_50_s of about 1.5 µM caffeine and a signal to noise ratio (reporter expression with 100 µM caffeine/ reporter expression without caffeine) of more than 50-fold (**Figure 5b; table S6**).

**Figure 5:**
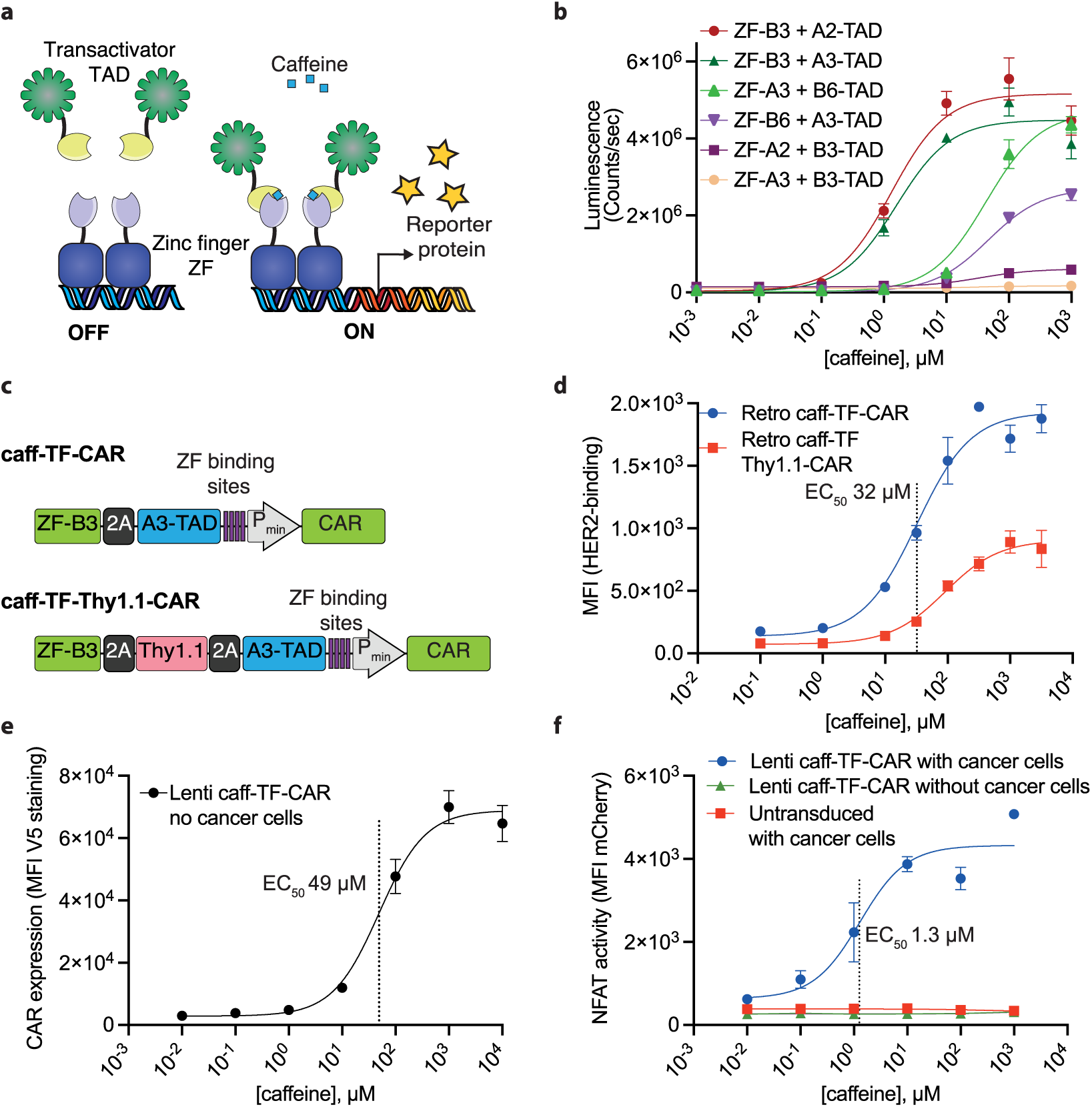
Characterization of Caffeine Responsive Zinc Finger Transcription Factors (ZF-TF). a, Schematic showing caffeine induced assembly of zinc finger proteins and transactivators, which can be functional (left) or nonfunctional (right). **b,** Dose response of heterodimeric caffeine inducible ZF-TFs with transient transfection in HEK293T cells. **c,** all-in-one caff-TF vectors for caffeine induced gene expression. **d,** Jurkat cells transduced with retroviral all-in-one caff-TF vectors for caffeine induced gene expression of a murine CAR. **e,** Jurkat cells transduced with the lentiviral all-in-one caff-TF vector for caffeine induced gene expression of a human CAR (corresponding to c, top). **e,** Jurkat cells transduced with the lentiviral all-in-one caff-TF vector for caffeine induced gene expression of a human HER-2 CAR with or without SKOV3 coculture, compared to untransduced Jurkat cells with SKOV3 coculture. Values in the dose response curves are the mean ± s.d. of at least n = 2 biological replicates.

Neither the reverse order of the same caffeine heterodimers in the transcription factors (ZF-A2 or A3 together with B3-TAD) nor the respective homodimers or the homodimer based on Humanized-4 resulted in effective caff-TFs with either absent or minimal fold changes and very low reporter gene expression (**Figure S4; table S6**). These results might be explained by the proximity of the caffeine nanobodies fused to DNA binding domains when bound to repetitive promoter elements. High local nanobody concentrations might favor the formation of inactive homodimers and block the binding of correctly assembled transcription factors. This hypothesis is consistent with the results that in the GEMS, the nanobody B3 had the lowest sensitivity for caffeine induced homodimerization. However, for the pairs ZF-B6 + A3-TAD or ZF-A3 + B6-TAD, we did not observe the same trend but greatly reduced caffeine sensitivity (**Figure 5b, d; table S6**).

The nonresponsive caff-TFs show that caffeine does not directly induce transcription from the reporter plasmid, and the control transcription factor ZF-TAD without nanobody fusions was not affected by any caffeine concentration, including 1000 µM. These results support that caffeine, even at very high concentrations, is not toxic to HEK293T cells.

Next, we tested the pair ZF-B3 + A3-TAD in an all-in-one retroviral vector containing the transcription factor parts linked with a P2A site and the inducible promoter elements for driving murine CAR expression. To test if this construct causes background signaling based on P2A readthrough resulting in ZF-B3-A3-TAD fusion proteins, we also generated a vector containing a Thy1.1 protein flanked by P2A sites separating the two parts of the split transcription factor (**Figure 5c**). We tested these vectors in Jurkat cells and observed dose-dependent caffeine inducible murine CAR expression with a signal to noise ratio of about 10-fold for both, suggesting that potential 2A readthrough is not causing background signaling (**Figure 5d**). However, the vector without Thy1.1 had an EC_50_ of about 30 µM, while the one with Thy1.1 had an EC_50_ of about 90 µM.

We chose the same caff-TF-CAR design without Thy1.1 to generate an all-in-one lentiviral vector for caffeine inducible human CAR expression. We tested this construct in a Jurkat cell line engineered to expresses mCherry in response to NFAT activation, which is a major signaling pathway downstream of CAR activation^34^. Consistent with the retroviral system, we observed caffeine-inducible CAR expression with an about 20-fold signal-to-noise ratio and an EC50 of about 50 µM.

To test for CAR activation, we co-cultured these engineered Jurkat cells with HER2-expressing SKOV3 cancer cells. NFAT signaling was strictly dependent on the presence of both caffeine and cancer cells, with a caffeine EC_50_ of about 1.3 µM. This result suggests that even low levels of CAR expression are sufficient to trigger downstream NFAT signaling. The EC_50_ values fall within a physiologically relevant range, as a single cup of coffee typically yields plasma caffeine concentrations around 15 µM^22^.

Together, these results demonstrate that an all-in-one lentiviral system, composed entirely of human or humanized components, enables tightly regulated, caffeine-inducible expression of functional CARs.

## Discussion

Tunable control over cell signaling or gene expression is of great interest for increasing the efficacy and safety of cell therapy^1,9,35^. In this work, we developed humanized caffeine-inducible homodimers and heterodimers to control cell function with caffeine. These tools are likely more suitable for clinical applications compared to previous systems that rely on nonhuman protein domains or use small molecule inducers with considerable side effects^11^. Nonhuman protein domains, such as receptor systems using viral TEV protease^35–37^ or fusing bacterial DNA binding proteins like dCas9 or TetR to viral transcriptional activators^14,38^, carry the risk of causing immune reactions against the equipped cells, likely reducing cell persistence and their efficacy in cell therapy^7–9^. Additionally, many previous systems are constrained by small molecules with less favorable pharmacokinetics, safety, and side effect profiles compared to the non-toxic and easily accessible molecule caffeine.

We demonstrate two key use cases of the engineered nanobodies by generating caffeine-inducible systems for cytokine signaling and synthetic transcriptional control, both using exclusively human or humanized protein domains. In parallel, Sylvander *et al*. independently developed a related heterodimerization system for caffeine-responsive split CARs, as described in the accompanying study. Our findings therefore suggest that the humanized heterodimerizing and homodimerizing nanobodies generated here will be broadly useful as modular components in controllable systems for cell therapy applications

The previous caffeine-responsive GEMS system relied on camelid nanobodies and murine EpoR domains^17^. Here, we humanized the nanobody and replaced the murine receptor domains with their human counterparts, achieving a humanized receptor for caffeine-induced STAT3 activation. As STAT3 signaling is one of the main pathways investigated for its role in counteracting T-cell exhaustion^39^, we constructed a lentiviral vector containing caff-GEMS and a STAT3-dependent reporter and confirmed its function in Jurkat T cells. In addition to direct effects derived from inducible STAT3 activity, the reporter can presumably be replaced with therapeutic cargo for caffeine-induced expression.

We used the humanized caffeine-induced heterodimers in a platform of human protein-based synthetic zinc finger proteins (ZFs) that bind a DNA sequence not present in the human genome^16^. These systems likely function without disturbing the native cell machinery, but as pointed out by the authors, face some challenges for small molecule-induced transcriptional control, as they were functionalized either with viral (NS3) or plant (ABI/PYL) derived protein domains or were fused to human estrogen receptor ERT2 for control with the drug tamoxifen and its active metabolite afimoxifene. Tamoxifen has several contraindications and associated risks, including being listed as a carcinogen that increases risks of endometrial cancer. Additionally, the very long half-life of 7 days for tamoxifen and 14 days for some active metabolites limits its versatility when used as a small molecule inducer for synthetic systems^13^. Therefore, the humanized nanobody heterodimers that sense caffeine, as a nontoxic inducer molecule with an average serum half-life of 1.5 - 9.5 h^22^, synergize favorably with the ZFs to bring this technology closer to clinical application.

Humanization was complicated by the fact that interface mutations were likely to change dimerization behavior of the nanobodies, and consequently all the first humanized sequences exhibited substantial background dimerization in the absence of caffeine. We therefore tested a few additional variants and assume that the additional humanizing interface mutation M61A was critical for reducing background dimerization by reducing interface contacts. The optimized humanized homodimeric and heterodimeric nanobodies now contain 6 - 7 amino acid differences to a human sequence outside of the CDRs. For comparison, the approved humanized biologic Caplacizumab^33^ contains 11 amino acid differences by the same analysis method. Therefore, caff-TFs and caff-GEMS have the potential to be compatible with clinical use.

We generated caff-TFs by fusing one of the caffeine binders to the ZF, and one to the human transactivator domain of RELA/p65. Heterodimerization of these two domains thus reconstitutes a functional transcription factor. Notably, heterodimerization in this context was critical and homodimerization did not result in functional caff-TFs. We speculate that ZF binding to the repeats in the synthetic promoter positions the caffeine nanobodies next to each other, resulting in high local nanobody concentrations that favor the formation of inactive homodimers. These likely bind DNA with high avidity while blocking the binding of correctly assembled TFs and thus reduce system performance. This hypothesis is supported by the observation that the best performing system contained nanobodies on the ZFs that had the lowest propensity of homodimerization. As the ZFs, the caffeine nanobodies, and the transcriptional activation domain (TAD) of RELA/p65 are each relatively small, the combination makes for a total of less than 2000 bp. Therefore, they fit into a single lentiviral or retroviral vector in addition to a minimal promoter that drives CAR expression. The observed tunable, caffeine-induced CAR expression in Jurkat cells demonstrates the suitability of this system for controlling therapeutic cargo.

In this work, we developed humanized systems for caffeine-inducible gene expression via cytokine signaling or direct transcriptional control. While clinical translation to improve the precision and safety of cell therapies will require further optimization, these systems provide a proof of concept for externally controllable, modular, tunable, and human-compatible tools with broad utility in engineered cells.

## Methods

### Computational Design of Caffeine-Induced Heterodimers

To generate protein modules that specifically heterodimerize based on the preexisting caffeine-controlled VHH homodimer structure (PDB id :6qtl), we used a positive/negative design approach^40^ to create a protein-protein interface that favors heterodimers while disfavoring homodimerization. We developed an algorithm, based on the existing COMETS (Constrained Optimization of Multistate Energies by Tree Search) method^41^ to achieve this. COMETS is a protein design tool that optimizes sequences for properties such as binding affinity, specificity, and stability across multiple protein states. The method combines a branch-and-bound approach using the A* search algorithm to efficiently find optimal sequences without exhaustive searching, providing provably optimal solutions under specified constraints and accommodating both discrete and continuous flexibility.

Our modified algorithm, variational COMETS (varCOMETS), improves upon COMETS by reducing the number of solutions explored in the highly rugged multistate optimization space in three ways: it first utilizes a belief propagation algorithm^42,43^ to efficiently compute tight upper bounds on the energies of negative states (e.g. the homodimers). Additionally, it incorporates the edge message passing linear programming algorithm^44^ to rapidly compute lower bounds on the energies of positive states (e.g., the heterodimers). Finally, varCOMETS introduces a dynamic node expansion strategy in the A* algorithm, reducing the number of possibilities explored, as presented in^45^. Overall, varCOMETS efficiently navigates the highly complex, multi-state protein design sequence energy landscape.

A total of 5 states were considered for the multistate design task, listed as follows. (**1**) The objective of the design was to optimize the synthetic heterodimer A-1:CFF:B-1. The negative design objective consisted of disfavoring the homodimers (**2**) A-1:CFF:A-1 and (**3**) B-1:CFF:B-1. At the same time, the design process favored the formation of the heterodimer over the monomeric forms of the designs. Therefore two additional negative states were added, (**4**) A-1:CFF and (**5**) B-1:CFF. COMETS and varCOMETS admit constraints during the search process to ensure that the predicted energies of monomers and complexes during the search process enforce the biophysical realism of the design. The following constraints were added: the energy of the heterodimer A-1:CFF:B-1 could exceed the energy of the wildtype VHH:CFF:VHH by at most 5 Rosetta Energy Units (REU)^46^, while each binary complex A-1:CFF and B-1:CFF could as well exceed the energy of the wildtype VHH:CFF binary complex by five REUs.

For the input, amino acid residues in the VHHs interacting with caffeine were discarded from consideration for the positive/negative design as we reasoned changes to these residues would likely disrupt the formation of the ternary complex between the regions of interest. As the complexity of the problem is high to fully explore, we divided the problem into three zones, Ap, Bp, and C here in each zone the method was run separately on a subset of residues. Zone Ap consisted of residues K45 and E91 in A-1 and K45 in B-1, which were allowed to mutate to any of {Q/N/R/E/D/K/S}, while residue 91 in B-1 was allowed to change rotameric conformation but not to mutate. Zone Bp consisted of A-1 residues Q41, R47, and B-1 residue D64, which were allowed to mutate to any of {Q/N/R/E/D/K}, while A-1 residues D64/K67 and B-1 residues Q41, R47 and K67 were allowed to change rotameric conformations but not mutate. Set C consisted of residues F49 and Y108 in both A-1 and B-1, which were allowed to mutate to residues {F/W/Y/L/I/A/V/H/G/D/Q/N/S/R/K/E}. A pairwise energy matrix between all modeled residues was obtained using the igdump program in the the Rosetta software suite^46^ and was used to compute energy matrices between all pairs of residues within each group. The top sequences all resulted in charged residues at the interface with opposite charges facing each other in the heterodimers and same charges facing each other in the homodimers. We chose the best seven heterodimer candidates based on the multistate Rosetta energy ((energy of the best conformation in the positive design for the complex - energy of the best conformation of each VHH in the unbound state) - ( energy of the best conformation for the first negative complex - energy of the best conformation for each VHH monomer of the first negative pair) - (energy of the best conformation for the second negative complex - energy of the best conformation for each VHH monomer of the second negative pair). The code is available under: https://github.com/LPDI-EPFL/caffeine-multi-state-design

#### Compound Preparation and Storage

Caffeine was prepared in Milli-Q water as 100 mM stocks and aliquots were stored at -20 °C.

#### Cell Culture

The Jurkat T cell line was cultured in RPMI-1640 (Gibco, 72400047). HEK293T cells were cultured in DMEM (Gibco, 10566016) for experiments and in RPMI-1640 for virus production. All medium was supplemented with heat-inactivated 10% FBS (Gibco, 26140079) and Pen/Strep (Gibco, 10378016). All cells were cultured at 37°C and 5% CO_2_ in a humidified incubator.

#### HEK293T Cell Experiments and Reporter Assays

For the HEK293T cell experiments and reporter gene assays, 16,000 cells per well were seeded with 100 µL of complete DMEM medium in the inner 60 wells of a 96-well plate, 24 h before transfection. The DNA/ polyethyleneimine (PEI; Polysciences Inc., 24765-1) ratio was 1:5. All plasmids are described in **Table S1**, all relevant proteins in **Table S2**. All annotated sequences can be downloaded under (https://benchling.com/leo/f_/edUxcK5h-caffeine-heterodimer/).

For GEMS, the plasmid mix for 30 wells was: GEMS vector – 3000ng ( or for heterodimers plasmid A - 1500 ng + GEMS plasmid B - 1500 ng), STAT3 (pLS392) expression plasmid – 500 ng, STAT3 reporter (pLS858)– 750 ng. For ZF-TFs, the plasmid mix for 30 wells was: ZF-nanobody – 1500 ng, nanobody-TAD – 1500 ng, ZF-reporter (pCK109) – 1000 ng.

The transfection mixture was vortexed, incubated for 20 minutes, and 50 µL/well was added to the cells. These numbers correspond to ∼130 ng DNA and 625 ng PEI per well. After 16 hours, the medium with the transfection mix was replaced with 100 µL per well of fresh complete medium containing the indicated drugs. After 24 h, supernatant samples were collected for quantification of the secreted reporter protein nanoluciferase. For the assay, 5 µL of cell supernatant was mixed with 5 µL of a nanoglo substrate:buffer solution (Promega, N1110) at a 1:50 ratio in a black 384-well plate and analyzed using a multiplate reader. For a more detailed protocol, please refer to ^47^.

#### Recombinant Lentivirus and Retrovirus Production and Transduction

24 h before transfection, 1.5 × 10^6^ HEK-293T cells/well were seeded in 2 ml of DMEM (10% FCS, 1% P/S) in a six-well tissue culture plate. For transfection, per well 700 ng of pVSV-G (VSV glycoprotein expression plasmid), 1.8 µg of delta8.2 (packaging plasmid for lentivirus) or pCL-Eco (packaging plasmid for retrovirus), and 4 µg of the lentiviral or retroviral vector and 20 µg PEI were used. After 16 h, the medium was exchanged for 1 mL RPMI (10% FCS, 1% P/S) and virus was produced for 48 h. For transduction, 3×10^5^ Jurkat cells were seeded into 12 well plates and mixed with the filtered (45 µm) virus-containing supernatant. After 16 h, the medium was exchanged for fresh RPMI and cells were expanded for another 5-12 days before the experiments.

#### Statistics

All calculations were performed using FlowJo (Version 10) and GraphPad Prism (Version 10). Exact p-values are shown in the figures where appropriate. The representative data from cell assays are presented as individual values overlayed with bars of mean values and error bars for s.d. or s.e.m. as indicated in the legend. For Jurkat and HEK293T cells, “n = 3” represents biological replicates, which are defined as different wells on the same plate. For all shown dose response curves, the EC_50_, and fold changes are shown in table S6.

#### CDR Annotations

We used the automatic CDR annotations provided by Benchling (Benchling [Biology Software]. (2024). Retrieved from https://benchling.com.). This is based on antibody numbering by the SAbPred/ANARCI and the IMGT CDR definitions^32^ and uses Sequence-based antibody CAnonical LOoP structure annotation (SCALOP)^48^.

#### Plasmid Construction

All plasmids are described in **Table S1**. Gene fragments (Twist Bioscience) were ordered with flanking regions overlapping with the target vector and inserted by Gibson assembly.

#### Flow Cytometry of Jurkat Cells

For human CAR or GEMS staining, we used a 1:100 dilution of an anti-V5 antibody (1:100, Invitrogen, 12-6796-42). For the murine CAR, we used Biotinylated Human Her2 (1:100, acrobiosystems, HE2-H822R-25ug) and PE-Streptavidin (1:100, BioLegend, 405203). The medium was exchanged for 20 µL/ well FACS buffer (PBS, 0.5 % BSA) containing the antibody or HER2 in round bottom 96 well plates and incubated for 30 min at 4 degrees. For HER2 staining, the protocol was repeated for incubation with streptavidin. Then, the cells were washed twice before measurement.

## Acknowledgements

The Swiss Cancer League (KFS-5032-02-2020) and the Swiss National Science Foundation (310030_197724, TMGC-3_213750) supported BC. The Strategic Focus Area “Personalized Health and Related Technologies (#2021-446)” of the ETH Domain and the Anniversary Foundation of Swiss Life for Public Health and Medical Research supported LS.

## Data Availability

All data is available in the main text or the supplementary information materials. Source Data are provided with this paper. Plasmids are available on request.

## Author Contributions

L.S., P.G., and B.E.C. developed the project. L.S., M.E., L.B., C.K., Y.G., P.D., J.S., T.E., S.G. designed and performed experiments and analyzed the data. P.G. developed and applied the varCOMETs algorithm to design the initial heterodimers. L.S. humanized the proteins. L.S. wrote the manuscript. All authors discussed findings and provided feedback on the manuscript.

