## supplementary material for "Humanized Caffeine-Inducible Systems for Controlling Cellular Functions"

| Plasmid table | |
|---|---|
| **Name** | **Description** |
| pCK028 | Heterodimer-B3 GEMS, SV40 promoter |
| pCK030 | Heterodimer-A2 GEMS, SV40 promoter |
| pCK032 | Humanized-5 GEMS, SV40 promoter |
| pCK069 | Heterodimer-B3 ZF, CMV promoter |
| pCK071 | Heterodimer-B3 TAD, CMV promoter |
| pCK072 | Heterodimer-A2 ZF, CMV promoter |
| pCK074 | Heterodimer-A2 TAD, CMV promoter |
| pCK079 | Heterodimer-B4 GEMS, SV40 promoter |
| pCK080 | Heterodimer-A3 GEMS, SV40 promoter |
| pCK084 | Heterodimer-B6 GEMS, SV40 promoter |
| pCK099 | Heterodimer-A3 ZF, CMV promoter |
| pCK101 | Heterodimer-A3 TAD, CMV promoter |
| pCK102 | Heterodimer-B6 ZF, CMV promoter |
| pCK104 | Heterodimer-B6 TAD, CMV promoter |
| pCK109 | 4 ZF binding sites, CMVmin, secreted nanoluciferase-Fc |
| pCK119 | Humanized-4 ZF fusion, CMV promoter |
| pCK120 | Humanized-4 TAD, CMV promoter |
| PLS103 | Design-1-R/ Design-2-R GEMS, SV40 promoter |
| pLS104 | Design-3-D GEMS, SV40 promoter |
| pLS105 | Design-3-R / Design-4-R GEMS, SV40 promoter |
| pLS106 | Design-4-D-D GEMS, SV40 promoter |
| pLS107 | Design-5-R-D GEMS, SV40 promoter |
| pLS1076 | Heterodimer-B2 GEMS, SV40 promoter |
| pLS1077 | Heterodimer-A1 GEMS, SV40 promoter |
| pLS1078 | Heterodimer-B1 GEMS, SV40 promoter |
| pLS108 | Design-6-E-D GEMS, SV40 promoter |
| pLS109 | Design-7-E-Q GEMS, SV40 promoter |
| pLS110 | Design-7-R GEMS, SV40 promoter |
| pLS188 | Design-1-E GEMS, SV40 promoter |
| pLS189 | Design-2-E-R GEMS, SV40 promoter |
| pLS190 | Design-5-E-R GEMS, SV40 promoter |
| pLS191 | Design-6-R-R GEMS, SV40 promoter |
| pLS392 | STAT3, SV40 |
| pLS410 | Camel WT nanobody GEMS, SV40 promoter |
| pLS630 | Humanized-1 GEMS, SV40 promoter |
| pLS671 | Humanized-2 GEMS, SV40 promoter |
| pLS674 | Humanized-3 GEMS, SV40 promoter |
| pLS785 | Humanized-4 GEMS, SV40 promoter |
| pLS858 | 4 STAT3 binding sites, CMVmin, nanoluciferase-Fc |

| | |
|---|---|
| pLS917 | Heterodimer-B5 GEMS, SV40 promoter |
| pLScar053 | All-in-one caff-TF retroviral vector, caff-TF-HER2-CAR, 4 ZF binding sites,<br>linker optimization and codon optimization |
| pLScar054 | All-in-one caff-TF retroviral vector, containing Thy1.1 |
| pLScar062 | All-in-one caff-GEMS lentiviral vector, caff-GEMS driving EGFP expression, 4 STAT3 binding sites |
| pLScar090 | All-in-one caff-TF lentiviral vector, caff-TF driving HER2-CAR expression, 4 ZF binding sites |

**Supplementary Table 1:** Description of all plasmids used. All annotated sequences can be downloaded under https://benchling.com/leo/f_/edUxcK5h-caffeine-heterodimer/

| Name | Description | Sequence |
|---|---|---|
| Camel WT, PDB:6qtl | WT camelid caffeine nanobody (homodimer) | QVQLVESGGGLVQAGGSLRLSCTASGRTGTIYSMAWFRQA<br>PGKEREFLATVGWSSGITYYMDSVKGRFTISRDKGKNTVY<br>LQMDSLKPEDTAVYYCTATRAYSVGYDYWGQGTQVTVSS |
| Design-3-D | Camelid caffeine nanobody, (heterodimer, R45D) | QVQLVESGGGLVQAGGSLRLSCTASGRTGTIYSMAWFRQA<br>PGKEDEFLATVGWSSGITYYMDSVKGRFTISRDKGKNTVY<br>LQMDSLKPEDTAVYYCTATRAYSVGYDYWGQGTQVTVSS |
| Design-3-R | Camelid caffeine nanobody, (heterodimer, D62R) | QVQLVESGGGLVQAGGSLRLSCTASGRTGTIYSMAWFRQA<br>PGKEREFLATVGWSSGITYYMRSVKGRFTISRDKGKNTVY<br>LQMDSLKPEDTAVYYCTATRAYSVGYDYWGQGTQVTVSS |
| Humanized-4 | Humanized caffeine nanobody (homodimer), best performer in GEMS | EVQLVESGGGLVQPGGSLRLSCAASGRTGTIYSMSWFRQA<br>PGKRREFLATVGWSSGITYYADSVKGRFTISRDNSKNTLY<br>LQMNSLRAEDTAVYYCTATRAWSVGYDYWGQGTLVTVSS |
| Heterodimer-A3 | Humanized caffeine nanobody (heterodimer, L48V, D62R), best performer in heterodimer GEMS and caff-TF | EVQLVESGGGLVQPGGSLRLSCAASGRTGTIYSMSWFRQA<br>PGKRREFVATVGWSSGITYYARSVKGRFTISRDNSKNTLY<br>LQMNSLRAEDTAVYYCTATRAWSVGYDYWGQGTLVTVSS |
| Heterodimer-B3 | Humanized caffeine nanobody (heterodimer, K43E, R44E, R45D, F47I, L48V), best performer in caff-TF | EVQLVESGGGLVQPGGSLRLSCAASGRTGTIYSMSWFRQA<br>PGEEDEIVATVGWSSGITYYADSVKGRFTISRDNSKNTLY<br>LQMNSLRAEDTAVYYCTATRAWSVGYDYWGQGTLVTVSS |
| Heterodimer-B6 | Humanized caffeine nanobody (heterodimer, K43E, R44E, F47I, L48V, R45D, E89R), best performer in heterodimer GEMS | EVQLVESGGGLVQPGGSLRLSCAASGRTGTIYSMSWFRQA<br>PGEEDEIVATVGWSSGITYYADSVKGRFTISRDNSKNTLY<br>LQMNSLRARDTAVYYCTATRAWSVGYDYWGQGTLVTVSS |
| EpoR ECD and TMD (F93A) | Human EpoR extracellular and transmembrane domain used in humanized GEMS. F93 renders the receptors insensitive to Epo. | SAPPPNLPDPKFESKAALLAARGPEELLCFTERLEDLVCF<br>WEEAASAGVGPGNYSFSYQLEDEPWKLCRLHQAPTARGAV<br>RFWCSLPTADTSSAVPLELRVTAASGAPRYHRVIHINEVV<br>LLDAPVGLVARLADESGHVVLRWLPPPETPMTSHIRYEVD<br>VSAGNGAGSVQRVEILEGRTECVLSNLRGRTRYTFAVRAR<br>MAEPSFGGFWSAWSEPVSLLTPSDLDPLILTLSLILVVIL<br>VLLTVLALLS |

**Supplementary table 2:** Relevant protein sequences. All annotated sequences can be downloaded under https://benchling.com/leo/f_/edUxcK5h-caffeine-heterodimer/

| Name | Mutations compared to the wildtype anti caffeine VHH | Plasmid name |
|---|---|---|
| Design-1-E (B) | K43E | pLS188 |
| Design-1-R (A) | K43R | PLS103 |
| Design-2-E-R (B) | K43E, E89R | pLS189 |
| Design-2-R (A)<br>(= Design-1-R) | K43R | PLS103 |

| | | |
|---|---|---|
| **Design-3-D (B)** | **R45D** | **pLS104** |
| **Design-3-R (A)** | **D62R** | **pLS105** |
| Design-4-D-D (B) | Q39D, R45D | pLS106 |
| Design-4-R (A) (= Design-3-R) | D62R | pLS105 |
| Design-5-R-D (B) | K43R, R45D | pLS107 |
| Design-5-E-R (A) | K43E, D62R | pLS190 |
| Design-6-E-D (B) | K43E, R45D | pLS108 |
| Design-6-R-R (A) | K43R, D62R | pLS191 |
| Design-7-E-Q (B) | R38E, E46Q | pLS109 |
| Design-7-R (A) | K43R | pLS110 |

**Supplementary table 3:** Mutations for caffeine induced heterodimers.

| **Name** | **Mutations compared to camel WT** | **Number of mutations compared to human (excluding CDRs)** | **Plasmid name** |
|---|---|---|---|
| Camel WT | - | 18 | pLS410 |
| Humanized-1 | A14P, T24A, E44G, R45L, K74N, G75S, V79L, D84N, K87R, P88A, Q114L | 7 | pLS630 |
| Humanized-2 | A14P, T24A, E44G, R45L, K74N, G75S, V79L, D84N, K87R, P88A, Y102W, Q114L | 7 | pLS671 |
| Humanized-3 | A14P, T24A, E44R, K74N, G75S, V79L, D84N, K87R, P88A, Y102W, Q114L | 9 | pLS674 |
| Humanized-4 | Q1E, A14P, T24A, A35S, E44R, M61A, K74N, G75S, V79L, D84N, K87R, P88A, Y102W, Q114L | 6 | pLS785 |
| Humanized-5 | Q1E, A14P, T24A, A35S, E44R, L48V, M61A, K74N, G75S, V79L, D84N, K87R, P88A, Y102W, Q114L | 5 | pCK032 |

**Supplementary table 4:** Mutations for humanized caffeine nanobodies tested in GEMS.

| **Name** | **Mutations compared to Humanized-4** | | | **Plasmid name** | | |
|---|---|---|---|---|---|---|
| | **Interface mutant** | **Heterodimerization mutant** | **Mutations compared to human** | **GEMS** | **ZF fusions** | **TAD fusions** |
| Humanized-4 | - | - | 6 | **pLS785** | pCK119 | pCK120 |
| Heterodimer-A1 | - | D62R | 7 | pLS1077 | | |
| Heterodimer-A2 | F47I, L48V | D62R | 6 | pCK030 | pCK072 | pCK074 |
| Heterodimer-A3 | L48V | D62R | 6 | **pCK080** | pCK099 | **pCK101** |
| Heterodimer-B1 | - | R44E, R45D | 6 | pLS1078 | | |
| Heterodimer-B2 | - | K43E, R44E, R45D | 7 | pLS1076 | | |
| Heterodimer-B3 | F47I, L48V | K43E, R44E, R45D | 7 | pCK028 | **pCK069** | pCK071 |
| Heterodimer-B4 | L48V | K43E, R44E, R45D | 7 | pCK079 | | |
| Heterodimer-B5 | L48V, A49S | K43E, R44E, R45D | 5 | pLS917 | | |

| | | | | | | |
|---|---|---|---|---|---|---|
| Heterodimer-B6 | F47I, L48V | K43E, R44E, R45D, E89R | 7 | **pCK084** | pCK102 | pCK104 |

**Supplementary table 5:** Mutations for humanized caffeine heterodimer nanobodies tested in GEMS and split zinc fingers. The plasmid names for the best performing pairs are marked in bold.

| **Figure** | **Name** | **Receptor plasmid** | **$EC_{50}$, [caffeine], µM** | **Reporter fold change [caffeine], 1000 µM/ [caffeine], 0 µM** | **Fold change in $EC_{50}$ homodimer/ heterodimer** |
|---|---|---|---|---|---|
| S1 | Design-1-E (A) | pLS188 | 0.73 | 20 | 8.8 |
| S1 | Design-1-R (B) | pLS103 | 0.064 | 3.3 | 0.77 |
| S1 | Design-2-E-R (A) | pLS189 | 0.12 | 7.4 | 1.5 |
| S1 | Design-2-R (B) (= Design-1-R) | pLS103 | 0.064 | 3.34 | 0.80 |
| **2d, S1** | **Design-3-D (A)** | **pLS104** | **1.1** | 21 | **4.7** |
| **2d, S1** | **Design-3-R (B)** | **pLS105** | **6.3** | 27 | **26** |
| S1 | Design-4-D-D (A) | pLS106 | 2.628 | 40 | 0.63 |
| S1 | Design-4-R (B) (= Design-3-R) | pLS105 | 6.314 | 27 | 1.5 |
| S1 | Design-5-R-D (A) | pLS107 | 24 | 23 | 21 |
| S1 | Design-5-E-R (B) | pLS190 | 3.9 | 32 | 3.5 |
| S1 | Design-6-E-D (A) | pLS108 | 0.94 | 9 | 3.3 |
| S1 | Design-6-R-R (B) | pLS191 | 0.38 | 9 | 1.3 |
| S1 | Design-7-E-Q (A) | pLS109 | 0.52 | 16 | 0.43 |
| S1 | Design-7-R (B) | pLS110 | 5.2 | 21 | 4.4 |
| S1 | Design-1-E + Design-1-R | pLS103 + pLS188 | 0.083 | 3.5 | A: 8.8<br>B: 0.77 |
| S1 | Design-2-E-R + Design-2-R | pLS103 + pLS189 | 0.079 | 3.5 | A: 1.5<br>B: 0.8 |
| **2d, S1** | **Design-3-D + Design-3-R** | **pLS104 + pLS105** | **0.24** | 10 | **A: 4.7**<br>**B: 26** |
| S1 | Design-4-D-D + Design-4-R | pLS106 + pLS105 | 4.2 | 31 | A: 0.63<br>B: 1.5 |
| S1 | Design-5-R-D + Design-5-E-R | pLS107 + pLS190 | 1.1 | 25 | A: 21<br>B: 3.5 |
| S1 | Design-6-E-D + Design-6-R-R | pLS108 + pLS191 | 0.29 | 15 | A: 3.3<br>B: 1.3 |
| S1 | Design-7-E-Q + Design-7-R | pLS109 + PLS110 | 1.2 | 27 | A: 0.43<br>B: 4.4 |
| 3a | Humanized 1 GEMS | pLS630 | - | 1.5 | - |

| | | | | | |
|---|---|---|---|---|---|
| 3a | Humanized 2 GEMS | pLS671 | - | 0.89 | - |
| 3a | Humanized 3 GEMS | pLS674 | - | 0.97 | - |
| **3a** | **Humanized 4 GEMS** | **pLS785** | **0.33** | **24** | **-** |
| 3a | Humanized 5 GEMS | pCK032 | 2.7 | 31 | - |
| 3e | Lentiviral caffeine GEMS | pLS410 | 0.1 | 5 | - |
| **4c** | **Heterodimer-A2 + B1** | **pCK030+ pLS1078** | **0.45** | **48** | A: 19<br>B: 7.5 |
| 4c | Heterodimer-A2 | pCK030 | 8.7 | 89 | 19 |
| 4c | Heterodimer-B1 | pLS1078 | 3.4 | 59 | 7.5 |
| **4d** | **Heterodimer-A3 + B1** | **pCK080+ pLS1078** | **0.43** | **27** | A: 19<br>B: 8.0 |
| 4d | Heterodimer-A3 | pCK080 | 8.2 | 79 | 19 |
| 4e | Heterodimer-A3 + B6 | pCK080 + pCk084 | 0.18 | 9.5 | A: 44<br>B: n.d. |
| 4e | Heterodimer-A3 (repeat) | pCK080 | 8.0 | 53 | 44 |
| 4e | Heterodimer-B6 | pCK084 | 111 | 38 | n.d. |
| 4f | Heterodimer-A2 + B3 | pCK080 + pCk028 | 0.0057 | 1.5 | A: 76<br>B: n.d. |
| 4f | Heterodimer-A2 (repeat) | pCK080 | 7.8 | 45 | 76 |
| **5b** | **ZF-B3 + A2-TAD** | **pCK069 + CK074** | **1.3** | **74** | - |
| 5b | ZF-A2 + B3-TAD | pCK072 + CK071 | 33 | 4 | - |
| **5b** | **ZF-B3 + A3-TAD** | **pCK069 + CK101** | **1.6** | **69** | - |
| 5b | ZF-A3 + B3-TAD | pCK099 + CK071 | - | 2 | - |
| 5b | ZF-A3 + B6-TAD | pCK099 + CK104 | 43 | 67 | - |
| 5b | ZF-B6 + A3-TAD | pCK102 + CK101 | 46 | 87 | - |
| 5d | Retro Caff-TF-CAR | pLScar053 | 32 | 15 | - |
| 5d | Retro Caff-TF-Thy1.1-CAR | pLScar054 | 90 | 7 | - |
| 5e | Lenti caff-TF-CAR | pLScar090 | 49 | 21 | |
| 5f | Lenti caff-TF-CAR with SKOV3 | pLScar090 | 1,3 | 6 | |

| | | | | | |
|---|---|---|---|---|---|
| 5f | Lenti caff-TF-CAR no cancer cells | pLScar090 | - | - | - |
| 5f | Untransduced Jurkat | - | - | - | - |
| S4b | ZF-TAD | pCK068 | - | 1 | - |
| S4b | ZF-B3 + B3-TAD | pCK069+pCK071 | - | 1 | - |
| S4b | ZF-A2 + A2-TAD | pCK072+pCK074 | - | 2 | - |
| S4b | ZF-A3 + A3-TAD | pCK099+pCK101 | 40 | 5 | - |
| S4b | ZF-B6 + B6-TAD | pCK102+pCK104 | - | 1 | - |
| S4b | ZF-Humanized 4 + Humanized 4-TAD | pCK119+pCK120 | 1.13 | 3 | - |

**Supplementary table 6:** Dose response curve analysis for representative figures. $EC_{50}$s and fold changes vary between experiments. Especially high fold changes (>20 fold) are strongly influenced by minor variations in background signaling. Best performers from each figure marked in bold.

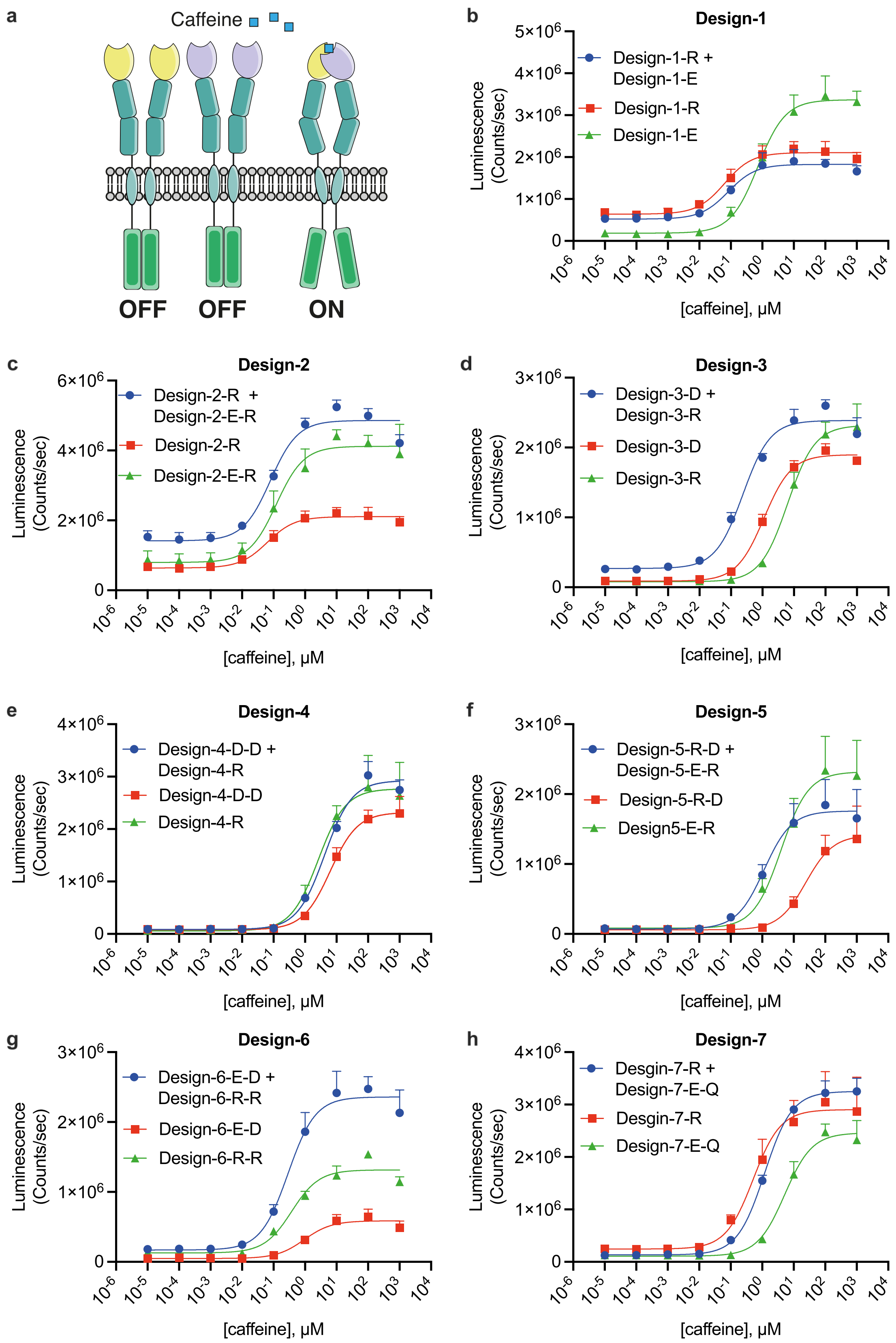


**Supplementary figure 1: Screening for caffeine induced heterodimerization.**
The plots show dose response curves of reporter protein expression in response to caffeine. Receptor sensitivity to caffeine induced heterodimerization was compared to the respective

homodimerization sensitivity of individual chains. Successful designs should show higher sensitivity to caffeine as heterodimers than as homodimers. Complex 3 shows this behavior. Some of the homodimers are shared between designs and the same homodimer data is used for clarity. Values are the mean ± s.d. of n = 3 biological replicates.

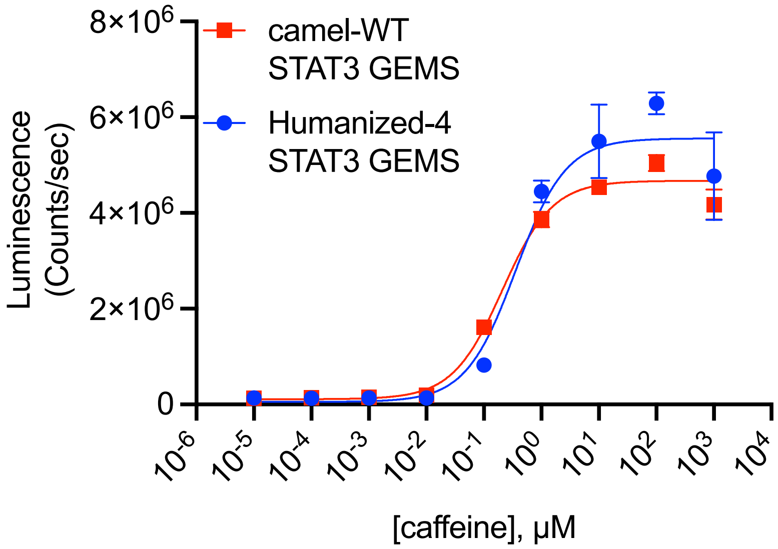


**Supplementary figure 2: Side by side comparison of Humanized-4 vs the original camelid nanobody.**
The plot shows dose response curves of reporter protein expression in response to caffeine. Values are the mean ± s.d. of n = 3 biological replicates.

```
Alignment Caplacizumab to anti-HIV-1 immunoglobulin heavy chain variable
region, partial [Homo sapiens]
Sequence ID: AXA20241.1 Length: 122
Range 1: 1 to 122

Identities:93/128(73%), Positives:100/128(78%), Gaps:6/128(4%)

Query  1    EVQLVESGGGLVQPGGSLRLSCAASGRTFSYNPMGWFRQAPGKGRELVAAISRTGGSTYY  60
            EVQLVESGGGLVQPGGSLRLSCAASG TFS   M W RQAPGKG E V+AIS +GGSTYY
Sbjct  1    EVQLVESGGGLVQPGGSLRLSCAASGFTFSSYAMSWVRQAPGKGLEWVSAISGSGGSTYY  60

Query  61   PDSVEGRFTISRDNAKRMVYLQMNSLRAEDTAVYYCAAAGVRAEDGRVRTLPSEYTFWGQ  120
             DSV+GRFTISRDN+K  +YLQMNSLRAEDTAVYYCA   V      V  +P  +  WGQ
Sbjct  61   ADSVKGRFTISRDNSKNTLYLQMNSLRAEDTAVYYCAKETV------VIAIPDAFDIWGQ  114

Query  121  GTQVTVSS  128
            GT VTVSS
Sbjct  115  GTMVTVSS  122
```

**Supplementary figure 3: Sequence alignment of Caplazizumab**
Sequence alignment of the approved humanized Caplacizumab[33] to a top ranked result from a blast search against human proteins The CDRs (IMGT annotation) are highlighted in yellow. Nonhuman residues and the human counterpart outside of the CDRs are shown in bold and underlined.

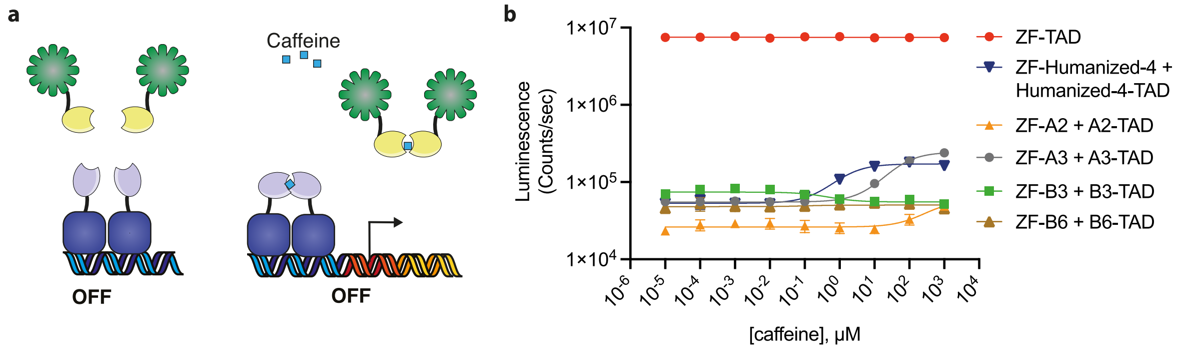


**Supplementary figure 4: Homodimeric caff-TFs**

**a,** schematic showing the presumed off state of the split transcription factors when neighboring nanobodies homodimerize. **b,** Dose response of homodimeric caffeine inducible ZF-TFs compared to the direct fusion of the ZF and the TAD as positive control. The Y-axis is on a log scale to compare the very high expression from the positive control with the homodimeric-nanobody based ZF-TFs. Values in the dose response curves are the mean ± s.d. of n = 3 biological replicates.
